# Highly quantitative measurement of differential protein-genome binding with PerCell chromatin sequencing

**DOI:** 10.1101/2024.07.12.603283

**Authors:** Alexi Tallan, Jack Kucinski, Benjamin Sunkel, Cenny Taslim, Stephanie LaHaye, Qi Liu, Jun Qi, Meng Wang, Genevieve C. Kendall, Benjamin Z. Stanton

**Affiliations:** Nationwide Children’s Hospital, Center for Childhood Cancer Research, Columbus, OH, USA; Molecular, Cellular, and Developmental Biology Program, The Ohio State University, Columbus, OH, USA; Cancer Biology Department, Dana-Farber Cancer Institute, Boston, MA, USA; Department of Pediatrics, The Ohio State University College of Medicine, Columbus, OH, USA; Department of Biological Chemistry & Pharmacology, The Ohio State University College of Medicine, Columbus, OH, USA

## Abstract

We report a universal strategy for 2D chromatin sequencing, to increase uniform data analyses and sharing across labs, and to facilitate highly quantitative comparisons across experimental conditions. Within our system, we provide wetlab and drylab tools for researchers to establish and analyze protein-genome binding data with PerCell ChIP-seq. Our methodology is virtually no cost and flexible, enabling rapid, quantitative, internally normalized chromatin sequencing to catalyze project development in a variety of systems, including in vivo zebrafish epigenomics and cancer cell epigenomics. While we highlight utility in these key areas, our methodology is flexible enough such that rapid comparisons of cellular spike-in versus non spike-in are possible, and generalizability to nuclease-based 2D chromatin sequencing would also be possible within the framework of our pipeline. Through the use of well-defined cellular ratios containing orthologous species’ chromatin, we enable cross-species comparative epigenomics and highly quantitative low-cost chromatin sequencing with utility across a range of disciplines.

## Introduction

Chromatin provides an interrelated network for structural and functional regulation of the DNA content in a eukaryotic cell. However, understanding a universal code with discrete combinations of histone modifications as keys to unlock chromatin has remained a central and meaningful challenge for our community. In order to understand the “keys” of chromatin regulation across the genome, our community would benefit from a low-cost, universal, and accessible process to generate, compare, and share chromatin sequencing data so that we can, together, define and understand the integrated patterns of the epigenome. This has motivated our current project – to provide an outward facing methodology encompassing both wetlab and drylab in order to enhance universal data generation, analysis, and sharing across teams.

The most well-studied and influential regulators of chromatin include: (**1**) post-translational modifications (PTMs) of the core canonical histone proteins around which DNA is wound (H2A, H2B, H3, and H4) as well as the non-canonical histones, which are also pervasive and subject to regulatory PTMs (including H1, H3.3, H2A.X, macroH2A, among others); (**2**) transcription factors (TFs), or proteins that alter rates of DNA transcription or replication via their specific chromatin interactions; (**3**) ATP-dependent chromatin remodelers, which frequently function as oncogenes in human tumorigenesis; (**4**) chromatin architectural machinery, including Cohesin complexes, CTCF, NIPBL, and WAPL which drive 3D chromatin structure; and (**5**) nucleic acid modifications, altering the structure-function relationships of DNA and RNA to direct chromatin mechanisms. A shared challenge across all of these chromatin regulatory modifications is how to quantitatively compare their abundance and localization across the genome, from one experiment to another, or even one lab to another. This is especially important if the experimental question entails deleting epigenetic regulators or causing global alterations in genome structure.

The advent and adoption of high throughput DNA/chromatin sequencing technologies, including ChIP-seq (Chromatin ImmunoPrecipitation followed by Sequencing)^1–4^, have become foundational and catalyzed a set of broadly popular methodologies. ChIP-seq can be applied to map histone modifications, transcription factor localization on chromatin, and indeed a myriad of other epigenetic events. The advantages of ChIP-seq include its reliance on largely non-enzymatic processes that avoid potential alterations of the regions of the epigenome that are sequenced (e.g., alternatives dependent on endonucleases cutting at open chromatin^5,6^). As a result, ChIP-seq remains highly useful even as a significant number of newer methods have been developed, which are fundamentally built upon the assay.

Despite this widespread use over many years, ChIP-seq as it is commonly performed has key limitations affecting its generalizability, data comparisons across teams of investigators, and indeed even across experiments in the same lab. These limitations have yet to be addressed in a highly quantitative and low-cost manner including both wetlab procedure and semi-automated data analysis. Chief among the limitations is our inability for highly quantitative ChIP-seq comparison across experimental conditions or cell lines. That is, while ChIP-seq can often be helpful in answering qualitative questions such as where a specific event occurs on chromatin (e.g., histone PTM, or TF binding) it is very difficult to draw confident conclusions in situations where a histone modification or TF binding event occurs at the same locus but in varying magnitudes. This information is often critical and varies depending on specific treatment conditions, timing within the cell cycle (where chromatin per cell may be altered), or across tumor cell lines of varying ploidy or genome sizes.

Additionally, as we progress towards an era of highly interdisciplinary science, epigenetics is often integrated with preclinical drug treatments (e.g., Entinostat, a clinical HDAC oncology candidate^7,8^; A485, a P300 preclinical oncology candidate^9^), many of which have potential to alter global histone modification states. If we are evaluating the basic epigenomic impact of preclinical molecules that target histone PTMs globally, there is a fundamental data processing challenge that arises and needs to be met. If changes occur everywhere on chromatin, then ironically, the changes are challenging to measure through next-generation sequencing compared to more specific, local epigenetic changes.

In attempts to address these fundamental limitations, methods have been introduced that include the addition of internal “spike-in” DNA content, measured by weight. This orthogonal DNA can be used to set a baseline across replicates and conditions, thereby enabling comparisons with greater clarity. However, some previous ChIP-seq protocols incorporating spike-in approaches were not designed for multicellular organisms^10^. Others used only highly divergent species’ DNA for spike-in^11^, or fixed amounts of spike-in chromatin^12^, or both^13^, creating technical hurdles related to inconsistent representation of the spike-in sequencing reads within datasets. While useful in providing quantitative comparisons in certain contexts, these methods also have drawbacks in that they can result in significant limitations in sequencing efficiency and the choice of antibody^11^, unnecessarily complicated procedures and the requirement for multiple antibodies^13^, or an inability to compare results across distinct genetic backgrounds^12,13^. This last point is especially significant given the high frequency of substantial changes in genomic content (aneuploidy) in both cancers and the immortalized cell lines often studied using ChIP-seq. Hence, we sought to establish a universal and low-cost strategy for data comparisons with PerCell ChIP-seq, which we describe herein.

## Advantages and Limitations of PerCell Chromatin Sequencing

Our approach takes advantage of “orthologous” genomes, meaning they can be resolved through next-generation sequencing, even when indexed the same way, within the same experimental sample. We have developed the orthologous cell spike-in to be within a framework of closely related genomes (e.g. human, mouse, zebrafish) and the mixing of experimental and spike-in cells at fixed ratios, within the same experimental sample, early in the workflow. The use of cells and not purified DNA for spike-in content enables the sensitive quantification and comparison of local and global differences in histone modification and transcription factor abundance/ localization across diverse treatment conditions and genomic contexts (**Figure 1**).

**Figure 1.**
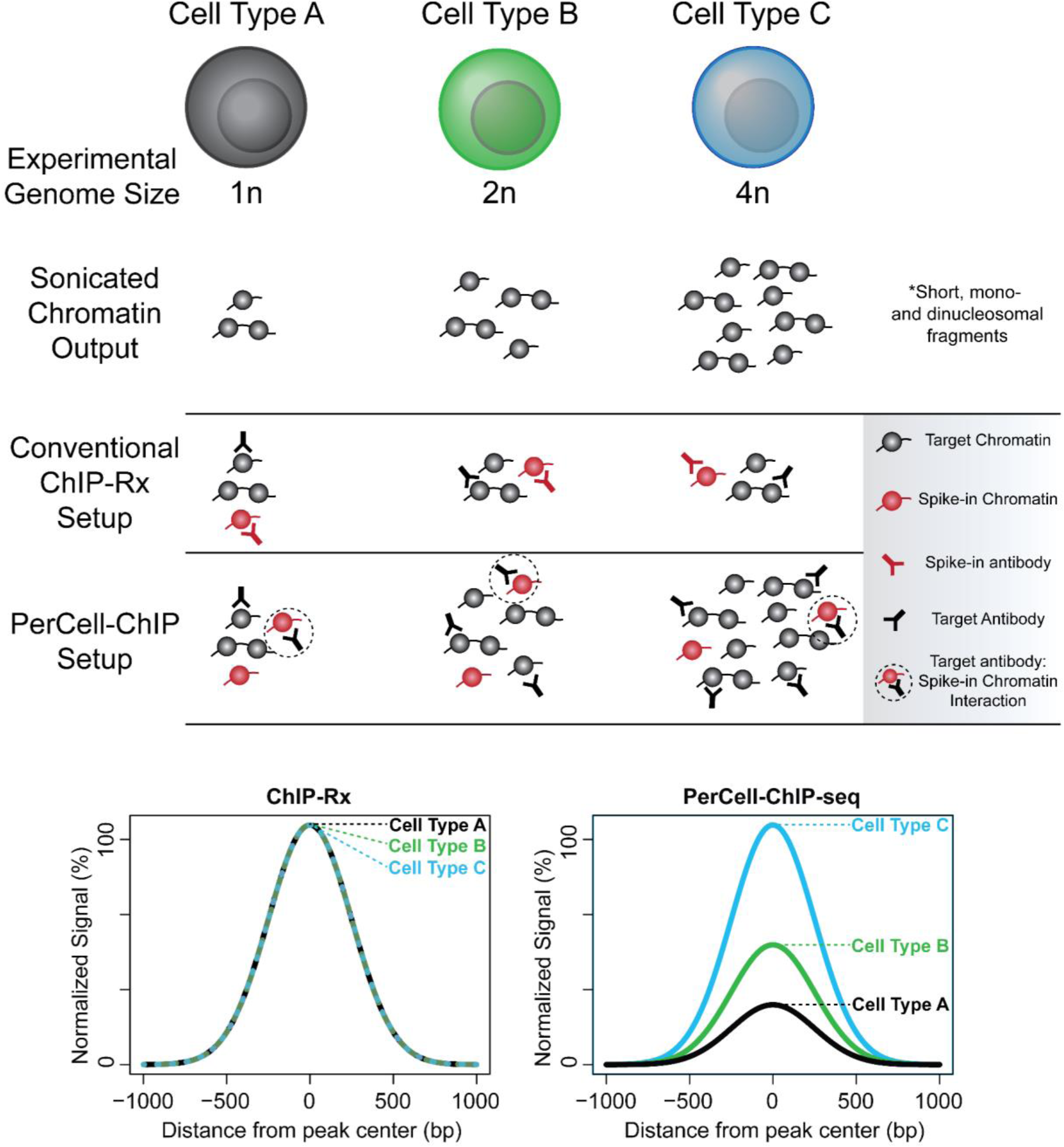
Rationale for PerCell normalized semi-automated ChIP-seq analysis. Conceptual representation and comparisons of ChIP results after analyzing cells of varying ploidy (1n, 2n, 4n) via conventional spike-in (ChIP-Rx) or PerCell normalization. PerCell methodology specifically enables the detection of changes in relative signal across cellular backgrounds with different chromatin content and the use of a single antibody.

The workflow is also facile for rapid data processing of standard, non spike-in ChIP-seq, which may be useful in cases where the ratio of spike-in versus test chromatin may be affected by PCR cycle number in library preparation, or by test cell types that generate very different sonication patterns from control spike-in cells. For example, human immune cells and mouse myoblast cells may be more challenging because the differentiation states may result in vastly different sensitivity to the mechanical shearing forces of sonication, but mouse fibroblasts and human myoblasts may be a technically more facile combination of experimental/spike-in cell chromatin based on comparative differentiation states and more similar distributions of chromatin sonication fragment lengths. Thus, consideration must be made for sonication efficiency across test and orthologous cell chromatin within an experimental setup, but the pipeline easily allows the user universal access to non-spike in comparisons of the same datasets (**Figure 2**).

**Figure 2.**
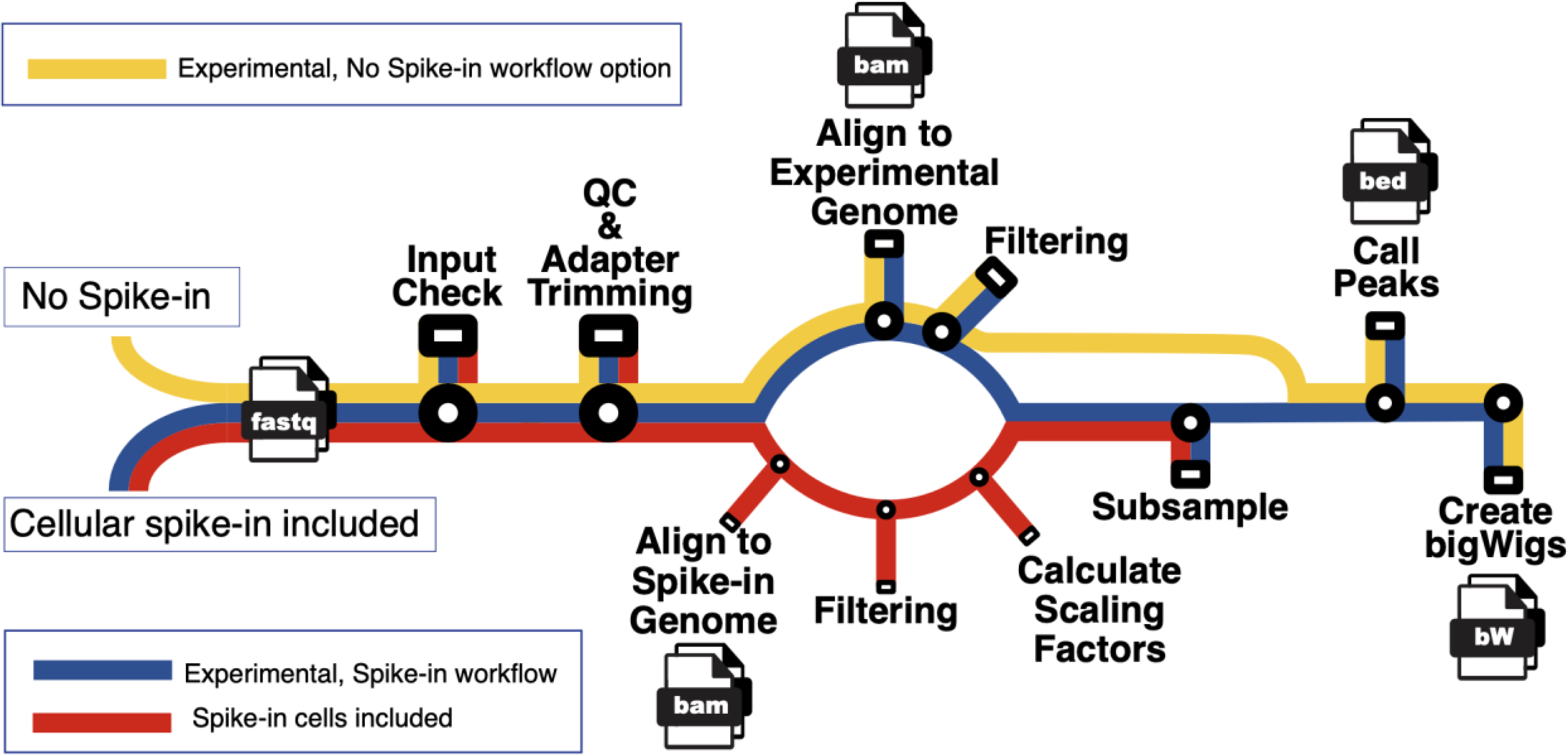
Subway map for PerCell workflow. Our workflow for semi-automated per cell normalization analysis pipeline with cellular spike-in (red), experimental chromatin (blue), or the workflow with no spike-in option (yellow). With a streamlined user interface, our pipeline will automate the central processes for cellular spike-in chromatin normalization per cell, including downsampling and peak calling to generate bigWig files. In our updated semi-automated per cell system, we include the option to either include or not include spike-in, which creates versatility and applicability to an increased range of researchers, and projects.

We have taken special consideration to benchmarking our method against previous approaches for chromatin or DNA spike-in. Specifically, in the context of highly quantitative comparisons between our methodological approach and previous approaches^11^, we have uncovered key differences in efficiency. In analyzing the number of reads aligned to the target (human) versus spike-in (mouse) genomes, we expect approximately 22.5% of mapped reads aligning to the spike-in genome, given a 3:1 human:mouse cellular ratio and accounting for the human genome being roughly 15% larger than the mouse genome. This is barring any effects on enrichment of one sample over another by the antibody used, and indeed we find fairly consistent percentages of 21-34% of input samples’ mapped reads aligning to the spike-in genome. For ChIP-ed samples, this range shifted slightly to 16-25%. This is in marked contrast to previous approaches^11^, where some ChIP-ed reads aligning to the spike-in (drosophila) genome equaled or even surpassed the number of reads aligning to the human genome. The relative percentages of spike-in versus experimental chromatin are important to consider, due to overall read depth and sensitivity within individual experiments as well as the potential for limitations on signal-to-noise within sequencing experiments.

## Applications and outlook

There is a diversity of applications for our methodology. These include, but are not limited to, comparisons of drug treatments, genetic deletions, and developmental changes in vivo. PerCell ChIP-seq enables efficient, cost-effective, non-commercial and reproducible strategies for epigenome sequencing. Our approaches do not rely on manufactured kits, and can be performed with common lab reagents, and thus provide benefits across experience and resource levels for researchers. Our method will generate rigorous maps of protein-genome binding and cis-chromatin interactions across cell states, and in vivo. *Further, there is no analytical pipeline intentionally designed for spike-in ChIP-seq analyses, which we will enable through these studies*.

Our pipeline is amenable to a range of spike-in setups, although we emphasize mouse-to-human and human-to-zebrafish quantitative epigenomics. Written using the Nextflow^14^ workflow management software, including customized implementations from the nf-core project^15^, our pipeline is highly scalable and reproducible across multiple execution platforms. With our workflows, comparisons are possible across treatment conditions, inhibition or functional disruption of global chromatin modifiers including targeted protein degradation, CRISPR/Cas9 knockout, and knockdowns. Additionally, our system has utility in human cancer cells with variable genomic content, and in a cell cycle context where the amount of chromatin per cell is no longer fixed, while the cell ratios remain constant (i.e. in S phase of the cell cycle).

While our pipeline was designed with PerCell ChIP-seq experiments in mind, its functionality is not limited to this protocol. The same normalization approach, and hence the same analytical processes, may be used for other assays involving spike-in chromatin, including the spike-in of fixed quantities of purified exogenous DNA as part of a standard ChIP-seq, Cut&Run^5^, or Cut&Tag^6^ assay. Together we anticipate that our PerCell experimental and semi-automated analysis workflows will be widely used to increase reproducibility across laboratories and across experimental conditions in cellular models and in vivo.

Our approach allows for the precise quantification of comparisons of global changes in the deposition of histone H3-acetylation after drug treatments in patient-derived rhabdomyosarcoma cells, with clinically promising molecules that either globally increase (entinostat, 0.5 μM) or globally decrease (A485, 0.5 μM) chromatin acetylation (**Figure 3**). Without PerCell measurements of differential histone H3 lys-27 acetylation, it becomes challenging to meaningfully interpret genome-wide data on the placement of this mark after either drug treatment (**Figure 3a,c,e**), whereas PerCell normalization enables direct quantitative comparisons (**Figure 3b,d,f**). When we examine the mapping of experimental and cellular spike-in reads from orthologous chromatin, we observe that there is less than 0.4% overlap between mouse/human mapped reads (**Supplementary Table S1**), meaning that the overwhelming majority of experimental or spike-in reads are mapped accurately to the correct genomes.

**Figure 3.**
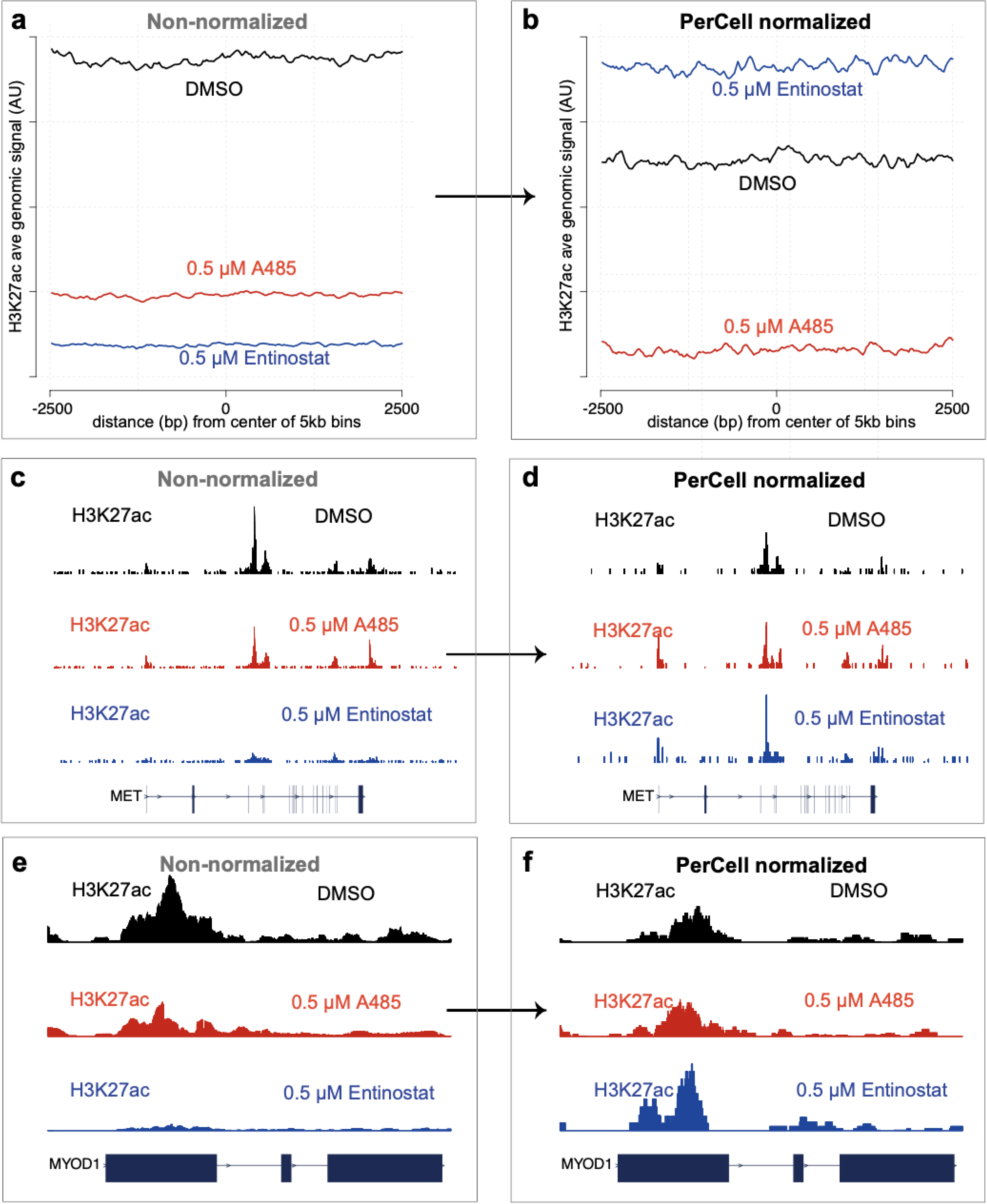
Data from human cancer cells (rhabdomyosarcoma, Rh4; with C2C12 mouse myoblast cellular spike-in), with anti-H3K27ac PerCell ChIPseq after DMSO treatment, A485 (0.5 μM), or Entinostat (0.5 μM) treatments. (a) Non-normalized and **(b)** PerCell normalized H3K27ac ChIPseq signal from Rh4 cells treated with DMSO, 0.5 μM A485 (histone acetyltransferase inhibitor), or 0.5 μM Entinostat (histone deacetylase inhibitor). Average signal is shown in arbitrary units (AU) across the entire genome divided into 5-kilobase (kb) bins. **(c,e)** Non-normalized and **(d,f)** PerCell normalized signal of the same samples at representative loci decorated with histone H3-acetylation, including MET **(c,d)** and MYOD1 **(e,f)**, respectively.

Of note, we report a specific application of our PerCell method to precisely map the genome-wide binding of a transcription factor (TF) chimera occurring in a rare childhood sarcoma (see **Procedure**, and **Figure 4**), for which we have developed new, rapid, and in vivo expression systems in zebrafish rhabdomyosarcoma models (**Figure 4a,b**), based on earlier contributions in this area^16^. Notably, when we rapidly express the TF-chimera PAX3::FOXO1 in vivo, PerCell normalization of anti-FOXO1 ChIP-seq^17,18^ reveals quantitative changes in the induction of H3K27ac at key target genes and binding sites (**Figure 4,c,d**). This is especially notable given that (1) these binding sites normally lack H3K27ac and (2) enhancer activation is typically driven by pluripotency transcription factors during these early embryonic zebrafish timepoints^19,20^. However, as we note in our companion manuscript (Kucinski et al, 2024), PAX3::FOXO1 binding also results in a quantifiable redistribution and therefore relative loss of H3K27ac at various loci as detected via PerCell ChIP-seq. Excitingly, our approach enables in vivo genome-wide binding measurements for PAX3::FOXO1 within hours after its expression, revealing chromatin state changes at the *her3* and *tfap2b* loci (**Figure 4e,f**). Excitingly, here the PerCell methodology is even more accurate and efficient in zebrafish/human systems than in human/mouse systems, and we find only approximately ∼0.001% of mapped reads overlapping between zebrafish/human genomes (**Supplementary Table S2, Supplementary Figure S1**). To our knowledge, this is the first reported in vivo genome-wide binding dataset for PAX3::FOXO1, and the initial report of highly quantitative differential comparisons of chromatin modifications states in vivo within hours of PAX3::FOXO1 induction. We anticipate further applications of PerCell methodology to measure TFs and TF-chimeras in human cancers, and their responsiveness to the activities of clinical or preclinical molecules that function through altering chromatin modification states.

**Figure 4.**
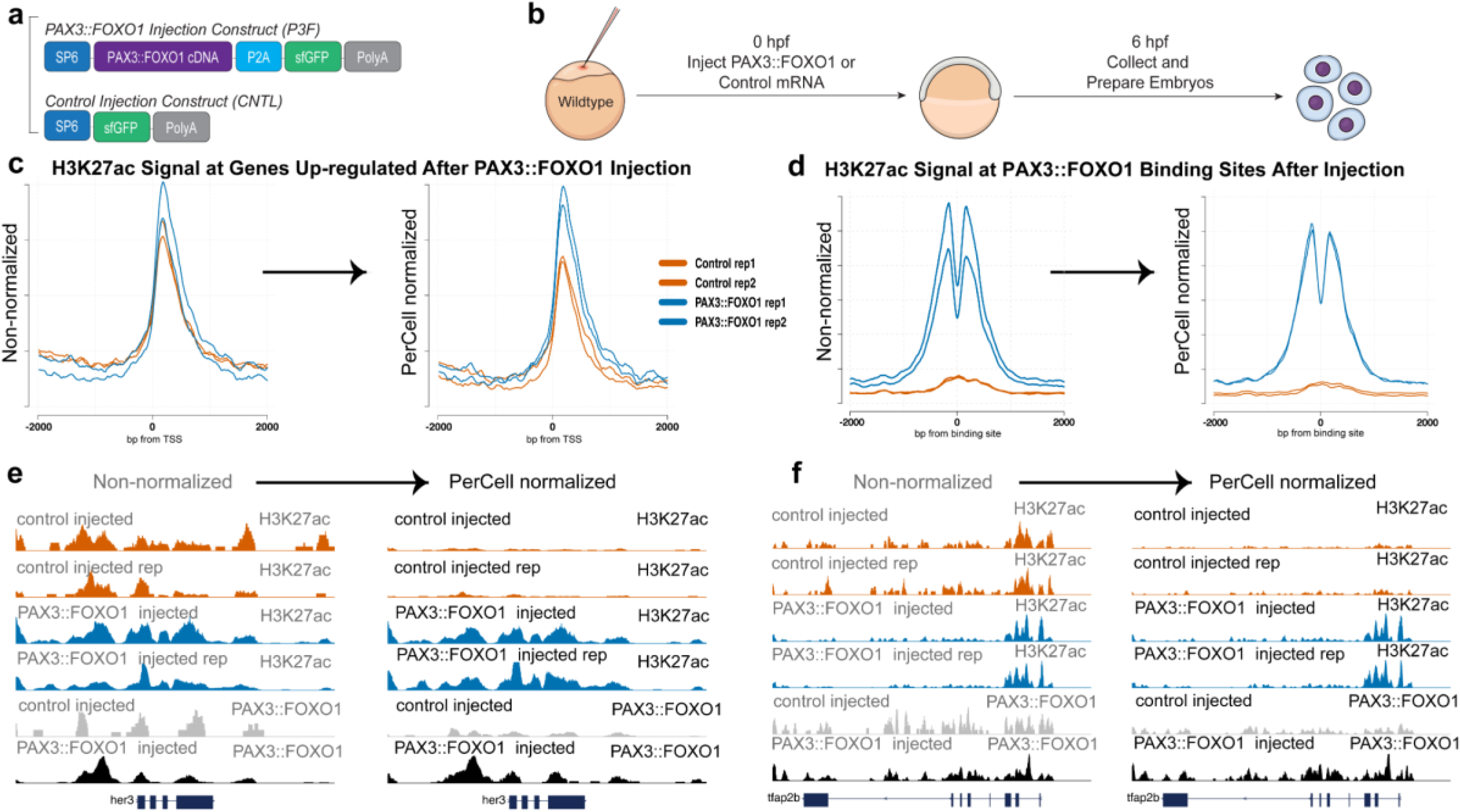
Zebrafish injected with Control (GFP) vs. PAX3::FOXO1 (P3F) mRNA, anti-H3K27ac and anti-FOXO1 (P3F) PerCell ChIPseq experiments. (a) Construct design is shown for PAX3::FOXO1 and control (GFP) mRNA with **(b)** our injection strategy in Zebrafish embryos, including experimental design workflow. **(c)** Profile plots are shown of non-normalized (left) versus normalized data for Control (GFP) versus P3F-injected at *up-regulated* genes or **(d)** known PAX3::FOXO1 (P3F) genomic binding sites containing DNA motifs comprised of either paired-domain/homeodomain (composite) or homeodomain motifs. Genome browser views are shown **(e)** at the her3 gene locus and **(f)** at the tfap2b locus, where P3F expression in zebrafish embryos increases local adjacent histone H3 lys-27ac levels, which can be measured with PerCell ChIPseq. In each case, the magnitudes of PAX3::FOXO1 binding are also measured with our PerCell workflow.

We also anticipate utility in the context of quantitative examination of protein-genome binding events with altered genome sizes, which are common in human cancers^21^. When we synthetically filter the apparent amount of cellular spike-in reads from experimental samples measuring CTCF binding, we are able to use PerCell normalization to obtain meaningful information for relative protein-genome binding with 25%, 50%, 75% and 100% of spike-in (mouse) reads retained (**Figure 5a,b**). With specific examination of the MYOD1 (**Figure 5c,d**) and FGFR4 (**Figure 5e,f**) loci, our PerCell workflow enables direct comparisons of CTCF binding levels with high resolution. There are key implications here, for contextualizing the loop extrusion model^22^, where examination not only of the convergence of CTCF sites bracketing loop domains, but also quantifying magnitudes of CTCF-genome binding could illuminate a central logic of chromatin folding. Thus, we illustrate utility of our approach in three distinct chromatin contexts, for architectural machinery (**Figure 5**), for the deposition of histone modifications and their drug sensitivities (**Figure 3**), and for cross-species genome-wide mapping and quantification of TF-chimeras that are penetrant drivers of childhood cancer (**Figure 4**). Further development of these workflows for defining altered protein-genome binding for TFs, other architectural regulators (e.g., Cohesin), a diversity of histone marks, and other epigenetic contexts, will be of high interest for researchers in the areas of pediatric oncology, epigenetics, and genomics.

**Figure 5.**
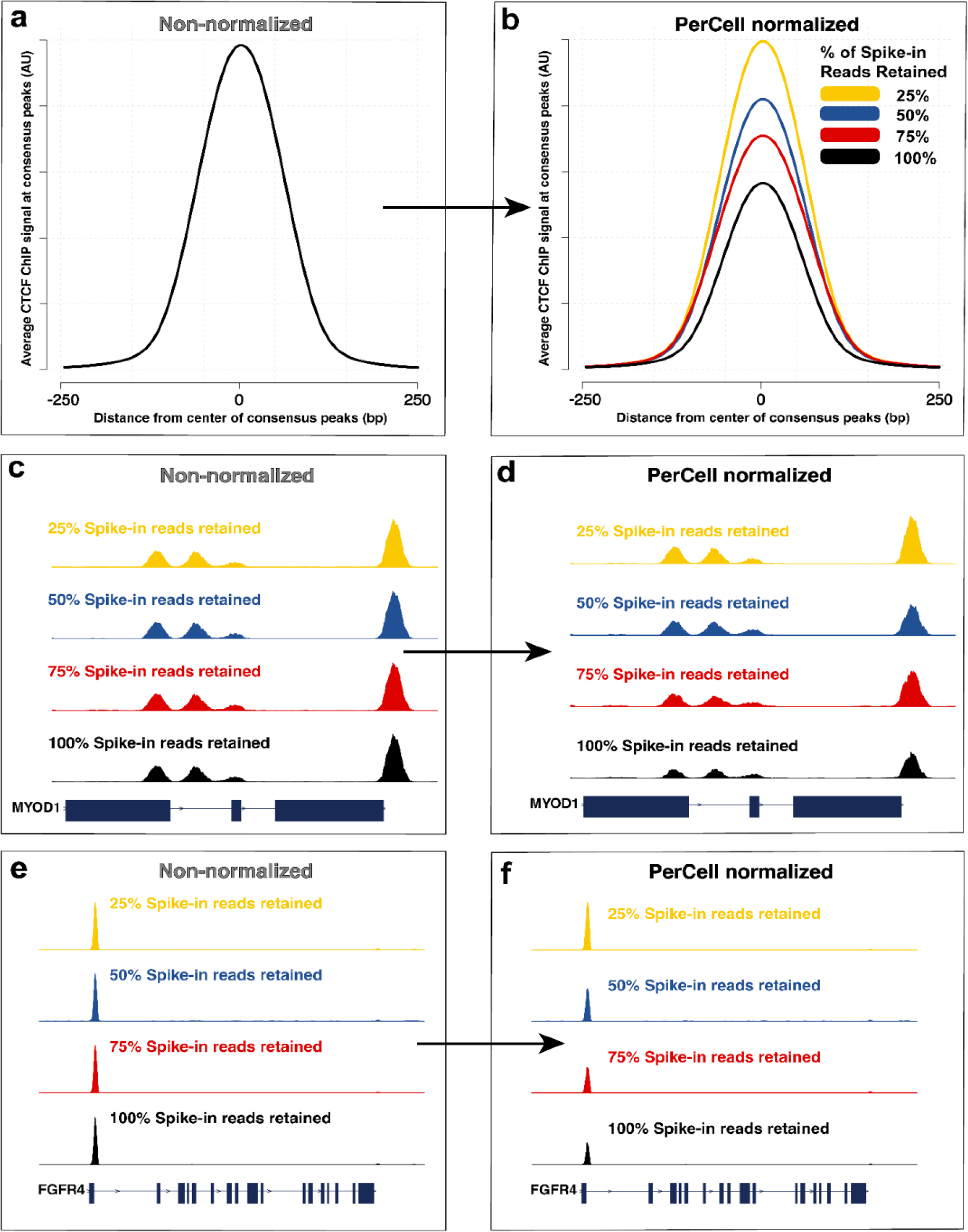
Quantitative PerCell differences in relative genome sizes. (a) Non-normalized and **(b)** PerCell normalized CTCF ChIP signal in Rh4 human rhabdomyosarcoma cancer cells, with varying percentages of cellular spike-in reads removed (resulting in 25%, 50%, 75%, or 100% of reads being retained) before application of the PerCell pipeline. Signal is shown +/− 250bp of consensus CTCF called peaks. **(c,e)** Non-normalized and **(d,f)** PerCell normalized signal at the MYOD1 **(c,d)** or FGFR4 **(e,f)** locus.

## Overview of Workflow

Our workflow is described in **Figure 2**, with a detailed overview of timing required per step in **Table S3** (see **Data setup and automated pipeline execution**). The required files for running our bioinformatic pipeline are fastq files, either gzip compressed or uncompressed. These files should contain sequencing reads for ChIP-seq or similar chromatin-based assays. While designed for handling mixed experimental and spike-in sequencing reads, the pipeline can also be run to skip all portions involving spike-in reads (e.g. for comparing spike-in normalization’s effects on analyses) and is thus suitable for analyzing data without spike-in included.

To start, the pipeline checks the formatting and existence of input files and then performs quality control analyses and any necessary trimming of sequencing reads to remove adapter sequences. All reads are next separately aligned to both an experimental and spike-in genome, with duplicates and poorly mapped reads being filtered out. Properly aligned spike-in reads are used to calculate scaling factors for all input and pulldown files. Reads aligned to the experimental genome are next subsampled using their respective scaling factors in order to normalize their signal based on a consistent inclusion of spiked-in chromatin/cells. Finally, downstream analyses on these normalized sequencing data are performed, including the calling of peaks (in bed file format) and generation of signal tracks (in bigWig format).

## Procedure

### Procedure 1: PerCell ChIP Experimental Procedure

#### Required Materials

##### Biological Materials

● WIK zebrafish, Zebrafish International Resource Center (ZIRC; https://zebrafish.org/)
  ○ NOTE: our methodology could be readily applicable to the study of an array of fusion oncoproteins and their early/immediate effects on chromatin modifications
  ○ NOTE: our approach could be generalizable to other cross-species approaches for mechanistic epigenomics
● RH30 cells (CRL-2061, ATCC)
● mRNA derived from Kucinski et al., 2024
● C2C12 (CRL-1772, ATCC)

##### Reagents

● DMSO, molecular biology grade cat#D8418, SIGMA
● Entinostat, cat#S1053, Selleckchem
  ○ NOTE: our protocol would be amenable to studying the early/immediate effects of a diversity of histone marks, including but not limited to H3 Lys27-acetylation
● A485, synthetic, Jun Qi lab, Dana-Farber Cancer Institute
  ○ NOTE: our protocol would be amenable to P300/CPB PROTACs as well
● Formaldehyde (methanol-free) cat#PI28906, Fisher Scientific
● Turbo DNase, cat#AM2238, ThermoFisher Sciencific
● Glycine, cat# G7126, SIGMA
● Pronase, cat#11459643001, SIGMA
● Proteinase K, recombinant PCR grade, cat#3115828001, SIGMA
● MinElute PCR Purification kit, cat#28006, QIAGEN
● Fetal Bovine Serum (10 x 50 mL), cat#A3160502, Fisher Scientific
● DMEM, cat#112-013-101CS, Quality Biological
● 2% E-Gel EX Agarose Gels, cat#G401002, Fisher Scientific
● Dynabeads Protein-A, cat#10002D, Fisher Scientific
● Dynabeads Protein G cat#10-003-D, Fisher Scientific
● T4 DNA Ligase, cat#M0202L, NEB
● End-Repair Enzyme mix (T4 DNA Pol + T4 PNK), cat#
● Klenow Fragment (3’→5’ exo-), cat# M0212S, NEB
● 100 mM dATP, cat#R0141, NEB
● 10 mM ATP, cat#10-297-018, Fisher Scientific
● 2.5 mM dNTP mix, cat#10-297-018, Fisher Scientific
● PE Adapter Oligo Mix (see PMID: 32051612 for sequences and preparation)
● Phusion Taq 2X Master Mix, cat#M0531S, NEB
● Illumina next generation sequencing index primers, PE1.0 (i5), and PE2.0 (i7), (see PMID: 32051612 for sequences)
● anti-FOXO1 Ab, C29H4, cat# 2880S, Cell Signaling
● anti-H3K27ac Ab, cat#39133, Active Motif
● Negative control ChIP-qPCR primers for zebrafish, cat#71035, Active motif
● Positive control ChIP-qPCR primers for zebrafish, nrp2a locus
  ○ FWD: CACAGCACTCATAAGCGAAGC
  ○ REV: TTGCACGGCGGTAAACAATC
● Negative control ChIP-qPCR primers for human RH30 cells, SOX18 locus
  ○ FWD: GCTCTTGGTTCTCTGTCCCT
  ○ REV: AGACAGACTGTGATGTGGGG
● Positive control ChIP-qPCR primers for human RH30 cells, FGFR4 locus
  ○ FWD: AAATTTGACCTTCGTCGGCAC
  ○ REV: CAGCTGTTGGCGATTTCACG
● Positive control ChIP-qPCR primers for human RH30 cells, QKI locus
  ○ FWD: TGCATGCTGGTGACAGATCA
  ○ REV: ACAGCGTCCTCTTTCAGCTT
● Positive control ChIP-qPCR primers for human RH30 cells, MYCN locus:
  ○ FWD: TCTCCAATTCTCGCCTTCAC
  ○ REV: GCGCTAACAGGTTTCTGTCC
● 100X SYBR Green, cat#S7563, Fisher Scientific
● AMPure XP beads, cat#A63880, Beckman Coulter
● Phusion High-Fidelity PCR Master Mix with HF Buffer, cat# M0531S, NEB
● NP-40/IGEPAL CA-630, cat# I3021-100ML, SIGMA
● Deoxycholate, cat#D6750-100G, SIGMA
● Triton X-100, cat#T8787-250ml, SIGMA
● Trypsin (sterile), cat#25200114, Fisher Scientific
● Protease Inhibitor Cocktail, cat#37491, Active Motif
● GlutaMax Supplement, cat#35050061, Fisher Scientific
● Water (molecular biology grade), cat#351-029-101CS, Quality Biological

##### Equipment

● Eppendorf™ Thermomixer, cat#RS232C, ThermoFisher
● Qubit Fluorometer, cat#Q33239, ThermoFisher
● EpiShear Probe Sonicator (110 V), cat# 53051, Active Motif (interchangeable with other sonication platforms)
  ○ NOTE: our protocol could be readily generalizable to Tn5-based and nuclease-based approaches for chromatin fragmentation as well
● EpiShear Cooled Sonication Platform (1.5 mL), cat# 53080, Active motif (interchangeable with other sonication platforms)
● E-Gel Power Snap Electrophoresis Device, cat# G8100, TheroFisher
● T100 Thermal Cycler, cat#1861096, BioRad
● LUNA Automated Cell Counter, cat#L10001, Logos Biosystem
  ○ NOTE: it is recommended to perform replicates of cell counting to ensure robustness of the PerCell approach

##### Buffers

● 0.5X Danieau’s buffer (29 mM NaCl, 0.35 mM KCl, 0.2 mM MgSO4·4H2O, 0.3 mM Ca(NO3)2·4H2O, 2.5 mM HEPES)
● Deyolking buffer (55 mM NaCl, 1.8 mM KCl, 1.25 mM NaHCO3)
● Phosphate Buffered Saline (PBS, sterile), cat#10010049, Fisher Scientific
● Embryo buffer 3 (5 mM NaCl, 0.17 mM KCl, 0.33 mM CaCl2, 0.33 mM MgSO4)
● TE buffer (pH 8.0), cat#351-011-721, Quality Biological
● 5M NaCl, molecular biology grade, cat# 351-036-101, Quality Biological
● 10% SDS, cat# 351-032-721, Quality Biological
● EB buffer, Qiagen, cat#19086
● 10% Triton X-100 (from cat#T8787-250ml, SIGMA, above)
● 0.1% sodium deoxycholate (from cat#D6750-100G, SIGMA, above)
● ChIP wash Buffer 1 (TE pH 8.0, with 0.1% SDS, 0.1% sodium deoxycholate, and 1% Triton X-100) (PMID: 24743988)
● ChIP wash Buffer 2 (TE pH 8.0, with 1% Triton X-100, 0.1% SDS, 0.1% sodium deoxycholate, and 200 mM NaCl) (PMID: 24743988)
● ChIP wash Buffer 3 (TE pH 8.0 with 250 mM LiCl, 0.5% NP-40, 0.5% sodium deoxycholate) (PMID: 24743988)
● End Repair buffer (330 mM Tris-acetate pH 7, 660 mM potassium acetate, 5 mM DTT)
● Klenow Buffer (NEBuffer 2), cat#B7002S, NEB
● 8M Lithium chloride (LiCl), cat# L7026-500ML, SIGMA
● T4 DNA Ligase Buffer (10X), cat#B0202S, NEB

### Sample Fixation

#### Timing: 1 hour (sample generation and collection will vary based on experimental set-up)

1) Collect experimental and spike-in samples/cells according to desired experimental condition and dissociate samples into single-cell suspension into Falcon or microcentrifuge tubes. Spin down cells at 300g for 5 minutes at room temperature.
  ● **CRITICAL STEP:** Experimental and spike-in samples MUST be from orthogonal species, so samples can be de-convoluted during genomic sequence alignment. Spike-in sample should be collected at the same time for all experimental conditions to minimize technical variation
2) Carefully remove supernatant and suspend cells in PBS and fix in 1% formaldehyde for 10 minutes at room temperature.
  ● Invert cells initially and every 2-3 minutes to ensure thorough and even fixation.
3) Quench fixation by addition of glycine solution to a final concentration of 125 mM and incubate on ice for 5 minutes.
  ● Invert cells initially and every 2-3 minutes to ensure thorough quenching.
4) Pellet fixed cells at 4°C, 1,200g for 5 minutes, carefully remove supernatant, and re-suspend in PBS with proteinase inhibitors.
5) Obtain cell count and aliquot desired cell amount into microcentrifuge tubes.
  ● **CRITICAL STEP:** Make sure cells are thoroughly suspended and not clumping to ensure accurate cell count. If cells remain clumpy following pipetting, fixation may not have been thorough.
  ● It is recommended to store cells in equal aliquots of 1-6 million cells, however aliquots can be later pooled to reach the desired cell number.
6) Pellet cell aliquots at 4°C, 1,200g for 5 minutes. Cell pellets can be stored for future use at –80 °C following snap freezing or immediately used.

### Chromatin Immunoprecipitation

#### Timing: 2-3 days

1) Thaw or continue with fixed cells. Combine experimental and spike-in cells in TE buffer (pH 8.0) supplemented with proteinase inhibitors. TE volume should be determined based on the recommended sonication volume according to the specific sonicator being used.
  ● **CRITICAL STEP:** The ratio of experimental to spike-in cells MUST be consistent across samples for comparative analysis as the pipeline determines the scaling ratio based on the consistent spike-in ratio in the samples.
  ● It is recommended to have ∼25% of reads be from the spike-in cells. However, this pipeline has successfully analyzed datasets with as high as ∼75% spike-in reads and as low as 2.5% spike-in reads.
  ● A higher percentage of spike-in cells may help serve as carrier chromatin to decrease the loss of the experimental sample throughout the protocol but will result in lower sequencing depth for your experimental samples.
2) Sonicate samples for the desired time and according to guidelines specific to the sonicator. Take 5 μL of sonicated DNA to serve as input DNA and to assess sonication efficiency. Store the remaining DNA at 4°C until sonication efficiency is confirmed.
  ● **CRITICAL STEP:** Sonicate the samples for the minimum amount of time required to achieve the desired fragmentation. For a histone mark ChIP-seq, DNA should be enriched around 200 bp. Meanwhile, a more transient factor (i.e. transcription factor or chromatin remodeler) ChIP-seq sample should be sonicated until the DNA is evenly distributed across fragment length to prevent over-sonication of the isotope.
  ● Experimental and spike-in cells, ideally, should sonicate at similar rates, which can be evaluated by sonicating both samples individually while determining the ideal sonication condition. However, experimental and spike-in cells MUST be sonicated together for this protocol to elevate sample variation.
3) Reverse DNA cross-link with one of the following reactions.
  a) 5 μL sonicated DNA, 8 μL 18.5 mg/mL Proteinase K, 4 μL 5M NaCl, and 90 μL H_2_O at 65°C with 400 rpm in a ThermoMixer for 1 hour
  b) 5 μL sonicated DNA, 20 μL TE buffer, 1 μL 10% SDS, and 1μL 18.5 mg/mL Proteinase K at 65°C overnight (less than 24 hours)
4) Purify DNA with a Qiagen MinElute PCR Purification kit (or comparable). Repeat the column binding step two times and incubate with 36 μL of warm EB buffer for four minutes to maximize DNA yield. Run 1 μL of purified input DNA on an E-gel (2%, EX) to confirm sonication efficiency.
  ● If samples are under-sonicated (i.e. DNA fragment is enriched at a greater length or is minimally enriched at 200) the remaining DNA can be further sonicated and steps 2-4 can be repeated.
  ● If samples are over-sonicated (i.e. fragment enrichment at less than 200 bp) chromatin immunoprecipitation should be repeated with new cells.
5) After successful sonication, adjust the remaining sonicated DNA (ChIP DNA) to a final buffer concentration of 1% Triton X-100, 0.1% SDS, 0.1% sodium deoxycholate, and 200 mM NaCl and centrifuge samples at 4°C, 17,949g for 10 minutes to spin-down any cellular debris.
6) Carefully, pipet the supernatant into a clean microcentrifuge tube and incubate with 2-4 ug of your desired primary antibody or according to the antibody manufacturer recommendation for 2 hours at 4°C with overhead rotation.
7) Take 40 μL per sample of Dynabeads Protein-A or Dynabeads Protein-G, according to the primary antibody isotype. Place beads on a magnetic rack to wash once in 200-400 μL of 1% Triton X-100, 0.1% SDS, 0.1% sodium deoxycholate, and 200 mM NaCl with brief overhead rotation. Re-suspend washed beads in the same volume initially taken of 1% Triton X-100, 0.1% SDS, 0.1% sodium deoxycholate, and 200 mM NaCl.
8) Add 40 μL of washed Dynabeads to protein samples and continue 4°C with overhead rotation overnight.
9) Wash immunoprecipitation with the various wash buffers below. Wash the immunoprecipitation by i) placing the microcentrifuge tube on the magnetic rack, ii) removing the supernatant, iii) adding 250-500 μL of the respective wash buffer, iv) briefly rotating until beads are re-suspended, and v) repeating
  ● Buffer 1 – 2 total washes of TE with 0.1% SDS, 0.1% sodium deoxycholate, and 1% Triton X-100
  ● Buffer 2 – 2 total washes of TE with 1% Triton X-100, 0.1% SDS, 0.1% sodium deoxycholate, and 200 mM NaCl
  ● Buffer 3 – 2 total washes of TE with 250 mM LiCl, 0.5% NP-40, 0.5% sodium deoxycholate
  ● Buffer 4 – 1 total wash with TE
10) Reverse DNA cross-link overnight by suspending beads in 100 μL TE, 2.5 μL 10% SDS, and 5 μL 18.5 mg/mL Proteinase K with overnight incubation at 65 °C (less than 24 hours).
11) Place beads on a magnetic rack, move supernatant to a clean microcentrifuge tube, and purify DNA with a Qiagen MinElute PCR Purification kit. Repeat the column binding step two times and incubate with 36 μL warm EB buffer for four minutes to maximize DNA yield.
12) <u>OPTIONAL:</u> Use qPCR to confirm immunoprecipitation effectiveness with 1 μL of ChIP and input DNA.
  ● qPCR can be done for regions in experimental or spike-in cells.

<u>STOPPING POINT</u>: Immunoprecipitated and input DNA can be stored at –20°C until the next step.

### Library Preparation

#### Timing: 1-2 days

1) Take 12-14 μL of ChIP DNA or 2-4 μL of input DNA for library preparation.
  ● **CRITICAL STEP:** It is recommended to prepare no more than 8 libraries simultaneously. If multiple days of library preparation are required, stagger replicates to account for potential batch effects.
2) Complete end-repair reaction according to the desired kit. Below is the response for the End-It DNA End-Repair Kit (Lucigen, ER81050) and incubate for 45 minutes at room temperature on a rocker.
  ● 34 μL DNA + H2O
  ● 5 μL 10X End Repair buffer (330 mM Tris-acetate pH 7, 660 mM potassium acetate, 5 mM DTT)
  ● 5 μL 2.5 mM dNTP
  ● 5 μL 10 mM ATP
  ● 1 μL End-Repair Enzyme mix (T4 DNA Pol + T4 PNK)
3) Purify DNA with a Qiagen MinElute PCR Purification kit and elute in 32 μL EB. Complete A-tailing reaction for 30 minutes on a ThermoMixer at 37°C, 300rpm
  ● 32 μL DNA (from End repair)
  ● 5 μL 10X Klenow Buffer (NEBuffer 2)
  ● 10 μL 1 mM dATP
  ● 3 μL 5 units/ μL Klenow Exo
4) Purify DNA with a Qiagen MinElute PCR Purification kit (or comparable) and elute in 2.2 μL EB. Complete linker ligation reaction with 30-60 minutes of room temperature incubation on a rocker.
  ● 22 μL DNA (from A-tailing)
  ● 3 μL 10X T4 DNA Ligase Buffer
  ● 2 μL PE Adapter Oligo Mix
  ● 3 μL T4 DNA Ligase (400 units/μL)
  ● **CRITICAL STEP:** Prepare PE Adapter Oligo mix, in advance, by diluting top and bottom adapter primers to a final concentration of 15 μM each. Heat the adapter mix at 95 °C for 5 min and cool at room temperature for 120 min.
  ● **STOPPING POINT:** Following incubation, samples can be stored at –20°C until the next step.
5) Select ligated products with AMPure beads (beads must be warmed to room temperature before selection). Add 30 μL (1.0X) AMPure beads, pipet well, and incubate at room temperature for 5 minutes and then on a magnetic rack for 5 minutes.
6) Remove supernatant and wash with fresh 80% ethanol on the magnetic rack, then air dry beads for 10 minutes to remove excess ethanol.
  ● **CRITICAL STEP**: The beads should be dried but not cracked. The exact incubation time may vary.
7) Add 23 μL EB to remove DNA from beads. Incubate at room temperature for 5 minutes and then on a magnetic rack for 5 minutes. Collect the supernatant.
  ● STOPPING POINT: Following collection, samples can be stored at –20°C until the next step.
8) Amplify DNA via SYBR-based qRT-PCR reaction. Only use ∼ half of the DNA as the other half can be amplified in case libraries are over-amplified or if more DNA is needed for sequencing following library preparation.
  ● Reaction:
    ○ 11.4 μL DNA
    ○ 12.5 μL Phusion Taq 2X Master Mix
    ○ 0.5 μL PCR primer lnPE1.0 (2X diluted) (i5)
    ○ 0.5 μL PCR primer lnPE2.0 (2X diluted) (i7)
    ○ 0.15 μL 100X SYBR Green
  ● PCR protocol:
    ○ 98°C for 30 seconds
    ○ Cycle – 98°C for 10 seconds, 65°C for 30 seconds, 72°C for 30 seconds
  ● **CRITICAL STEP:** Stop the reaction before it plateaus to minimize PCR duplicates. It is recommended to start with 10 cycles for input DNA or highly abundant histone marks/factors ChIPs and 15 cycles for less abundant histone marks/factors. Cycle number will vary by experiment.
  ● **CRITICAL STEP:** Ensure each sample has a distinct primer set to be de-convoluted following sequencing.
9) Purify amplified libraries using AMPure beads (beads must be warmed to room temperature before selection). Add 45 μL (1.8X) AMPure beads, pipet well, and incubate at room temperature for 5 minutes and then on a magnetic rack for 5 minutes.
10) Remove supernatant and wash with fresh 80% ethanol on the magnetic rack, then air dry beads for 10 minutes to remove excess ethanol.
  ● **CRITICAL STEP**: The beads should be dried but not cracked. The exact incubation time may vary.
11) Add 20 μL EB to remove DNA from beads. Incubate at room temperature for 5 minutes and then on a magnetic rack for 5 minutes. Collect the supernatant. Store DNA at –20°C.
12) Assess DNA libraries by 1) running on E-Gel or bioanalyzer to confirm library fragments are ∼ 300 bp and/or 2) obtaining DNA concentration by Qubit.

### Notes for ChIP-seq with zebrafish PAX3::FOXO1 mRNA injection model

Adult wildtype zebrafish (*Danio rerio*) were maintained in an aquatics facility in compliance with the Guide for the Care and Use of Laboratory Animals. However, various mutant zebrafish strains can be used to assess potential cooperativity in the model system. The human PAX3::FOXO1 coding sequence was synthesized and cloned from Shapiro et al., 1993 with the addition of *Pac1* and *Asc1* enzyme digestion sites. A PAX3::FOXO1-2A-sfGFP plasmid was generated for protein visualization by cloning the PAX3::FOXO1 product into pCS2+MCS-P2A-sfGFP (Addgene plasmid #74668)^23^. mRNA was generated by SP6 reaction of linearized PAX3::FOXO1-2A-sfGFP DNA with *Not1* digestion. Single-cell zebrafish embryos were injected with 100 ng/μL PAX3::FOXO1-2A-sfGFP a drop size diameter of 0.15 mm. A control mRNA construct was similarly injected with equal molarity. DNA constructs were generated by linearizing the pCS2+MCS-P2A-sfGFP plasmid with *Not1* digestion or amplifying out sfGFP and polyA tail from pCS2+MCS-P2A-sfGFP and transcribing DNA into mRNA as mentioned.

Following injection, zebrafish cells were incubated at 32°C for 5.25 hours to reach the proper developmental stage. Then, they were dechorionated with pronase, de-yolked in deyolking buffer (55 mM NaCl, 1.8 mM KCl, 1.25 mM NaHCO3), and washed with 0.5X Danieau’s buffer (29 mM NaCl, 0.35 mM KCl, 0.2 mM MgSO4·4H2O, 0.3 mM Ca(NO3)2·4H2O, 2.5 mM HEPES). Embryos were dissociated into single cells by thoroughly pipetting in dPBS. Samples were fixed as described in Procedure 1 and stored until enough cells were collected. ChIP-seq and library preparation was completed as described above. Generally, we could obtain approximately 1 million cells with 500 embryos, which took 1 hour of injecting. PAX3::FOXO1 ChIP-seq (4 μL anti-FOXO1 – Cell Signaling, C29H4) used 2 million zebrafish cells and 250,000 RH30 cells for spike-in. Immunoprecipitation was validated with qPCR using *SOX18* (negative) and *FGFR4* (positive) primers for RH30 cells and negative control (Active motif, 71035) and *nrp2a* (positive) primers for zebrafish cells. H3K27ac ChIP-seq (4 μL anti-H3K27ac – Active Motif, 39133) used 1 million zebrafish cells and 1 million RH30 cells for spike-in. Immunoprecipitation was validated with qPCR using *SOX18* (negative) and *QKI* or *MYCN* (positive) primers for RH30 cells.

### Procedure 2: Bioinformatic Analysis using PerCell Pipeline

#### Required Hardware

● Computer with Unix-based operating system
● A computer cluster or cloud-based computing platform for executing the pipeline (currently the pipeline is tested for compatibility with SLURM-managed platforms and AWS Batch service)

#### Required Software

Nextflow^14^ installation: https://www.nextflow.io/docs/latest/install.html

Singularity^24^ installation: https://docs.sylabs.io/guides/latest/admin-guide/installation.html

### Data setup and automated pipeline execution

#### Timing: 30 min-1 h setup; 2-8 h execution

**1.** Install both Nextflow and Singularity. Ensure both are properly running on the system.
**2.** If necessary, download files for chosen experimental and spike-in reference genomes in fasta format. Example commands for downloading and uncompressing human (hg38), mouse (mm10), zebrafish (danRer11), and fly (dm6) from UCSC can be found below:
  $ wget http://hgdownload.soe.ucsc.edu/goldenPath/hg38/bigZips/hg38.fa.gz
  $ gunzip hg38.fa.gz
  $ wget http://hgdownload.soe.ucsc.edu/goldenPath/mm10/bigZips/mm10.fa.gz
  $ gunzip mm10.fa.gz
  $ wget http://hgdownload.soe.ucsc.edu/goldenPath/danRer11/bigZips/danRer11.fa.gz
  $ gunzip danRer11.fa.gz
  $ wget http://hgdownload.soe.ucsc.edu/goldenPath/dm6/bigZips/dm6.fa.gz
  $ gunzip dm6.fa.gz
**3.** Create a csv file with names of samples, full locations of fastq files, antibodies/treatment used, and associated control sample names. Example input csv files are available at the PerCell GitHub repository (https://github.com/lextallan/PerCell).
  3.1. First line serves as column headings and must consist of the following: “sample,fastq_1,fastq_2,antibody,control”
  3.2. Each additional line follows that format for each sequenced sample
  3.3. Sample names should be unique, unless the same sample has been re-sequenced. Additional sequencing of the same sample can use identical sample names and the pipeline will merge the associated fastq files before alignment.
  3.4. Samples with multiple replicates should have sample names ending in a similar pattern followed by the number of the replicate, e.g. “_1”/ “_2” or “_rep1”/ “_rep2”. These will be treated as replicates by the pipeline for downstream analyses.
  3.5. fastq_1 and fastq_2 columns should contain the absolute path to the files, in an accessible directory. Files may be gzipped.
  3.6. Antibody column should contain info about the antibody used as well as any treatments done on the sample. Samples with identical antibody columns will be combined for peak calling. For example, samples created using antibody *A* but distinct cell lines/ treatments *B* or *C* could be written as “antibodyA_B” and “antibodyA_C”, respectively. For input samples, this column should be left blank.
  3.7. Control column should contain the <u>exact</u> name of the sample for the associated input, <u>including replicate information</u>. For example, “sampleA-Input_rep1” or “sampleA-Input_rep2” and NOT “sampleA-Input”.
**4.** OPTIONAL: Edit the file named nextflow.config with parameter choices (see execution step below for specific explanations) and options specific to the computing environment used (many organizations have suggested nextflow setup and config options, as collected by the nf-core community: https://nf-co.re/configs). The PerCell config file can be manually adjusted after directly downloading from our GitHub repository, (https://github.com/lextallan/PerCell/blob/master/nextflow.config) using the command
  $ git clone https://raw.githubusercontent.com/lextallan/PerCell/master/nextflow.config Alternatively, parameter options can be specified at the command line at the point of pipeline execution.
**5.** Execute the main pipeline script, PC.nf, with desired parameters (explained below). For ease of use, the entire pipeline and all potential parameters can be launched and customized from this single command. *Make careful note of whether one or two dashes are used in the following parameters. Single dashes are for general nextflow options, double dashes are for pipeline-specific parameter choices*.
  $ nextflow run lextallan/PerCell/PC.nf -profile singularity (tells nextflow to download/ use singularity images for each process) ––input: path to csv input file (see above for required format) ––outdir: path to directory where the pipeline’s output will be found ––aligner: choice of alignment tool with which samples will be aligned, <bowtie2,chromap> (default: bowtie2) ––experimental: species identify from which the experimental cells originate, <human,mouse,zebrafish,fly> (default: human) ––spikein: species identify from which the cells used for spike-in originate, <human,mouse,zebrafish,fly> (default: mouse) ––human_fa: path to human reference fasta file, if used as either experimental or spike-in genome ––mouse_fa: path to mouse reference fasta file, if used as either experimental or spike-in genome ––zebrafish_fa: path to zebrafish reference fasta file, if used as if used as either experimental or spike-in genome ––fly_fa: path to fly reference fasta file, if used as if used as either experimental or spike-in genome ––skip_fastqc: choice of whether or not to skip FastQC quality control assessments, <true,false> (default: false) ––skip_trimming: set to true if fastq files have already been trimmed of adapter sequences with an algorithm such as TrimGalore, <true,false> (default: false) ––save_trimmed: if set to true, trimmed fastq files will be saved as output, <true,false> (default: false) ––override_spikeinfail: the pipeline will not attempt to downsample experimental samples if spike-in content is detected to be below 0.5% of properly aligned total reads, setting this parameter to true will ignore this threshold and attempt normalization regardless of detected spike-in reads, <true,false> (default: false) ––skip_downsample: set to true to disable PerCell normalization and carry out a standard ChIP-seq pipeline analysis, <true,false> (default: false) ––seed: random seed used to downsample reads. default: 0 ––macs2_peak_method: choice of method with which to score macs2 bedgraphs for peak calling, <ppois,qpois,subtract,logFE,FE,logLR,slogLR,max> (default: ppois) ––macs2_cutoff: p/q-value cutoff used for calling peaks; unlike macs2’s default ‘callpeak’ command, cutoffs are taken in –log10(x) form. If using IDR, a particularly relaxed cutoff is highly recommended. For ChIPs targeting broader histone marks or when there are >2 replicates, IDR is not recommended and a more stringent cutoff of 1.301 (p/q-value of 0.05) or even 2 (p/q-value of 0.01) is suggested instead (default: 0.2218) ––macs2_bigwig_method: choice of method with which to generate bigWig tracks for visualizing ChIP data within a genome browser, <ppois,qpois,subtract,logFE,FE,logLR,slogLR,max> (default: ppois) ––skip_idr: if the IDR (Irreproducible Discovery Rate, https://github.com/nboley/idr) framework should be used to statistically identify significant peaks across two replicates. According to authors, the software is not designed for use with particularly broad peaks (e.g. those call in a ChIP for H3K9me3) and requires using a relaxed macs2 cutoff value in order to properly function, <true,false> (default: true) ––idr_cutoff: cutoff value over which peaks will not be included within the output (default: 0.05) ––skip_consensus: choice of whether consensus peaks should be identified across replicates, especially recommended along with a more stringent macs2 cutoff if there are >2 replicates, <true,false> (default: false) ––skip_annotation: whether or not to use HOMER’s ‘annotatePeaks’ tool for annotation of peaks with nearest genes and genomic features <true,false> (default: false) ––skip_motif: whether or not to use HOMER’s ‘findMotifsGenome’ tool to find enriched DNA sequence motifs in called peaks <true,false> (default: false)

## Anticipated Results

PerCell ChIP-seq assays and their analysis using our pipeline are expected to generate similar qualitative readouts of protein:DNA interactions or histone modification localization to conventional ChIP-seq workflows, while additionally providing relative quantifications for these data across distinct replicates, lab groups, cellular treatment conditions, and genetic backgrounds. We expect our spike-in based normalization method and pipeline to enable the user to identify global changes in transcription factor or histone modification abundance throughout the genome, while also accounting for differences in the chromatin states being investigated (e.g. due to aneuploidy). Additionally, the flexible choice of spike-in cells in a closely related species decreases barriers associated with requiring multiple antibodies or specifically prepared chromatin/ cell lines.

After initiation of the pipeline, the user specified input csv (“––input”) is first checked to ensure proper formatting and that all files both exist and are accessible to the pipeline. All pipeline outputs are stored in a directory named after the choice of alignment tool (“––aligner”) within the user-specified output directory (“––outidir”). When the applicable options are enabled (i.e. when “––skip_trimming” and “––skip_fastqc” set to false), adapter sequences are trimmed from reads and quality control data is collected and made available to the user via the MultiQC software^25^ in readable html format (*MultiQC-Report-for-PerCell-pipeline.html*). For user convenience, trimmed fastq files can be retained and organized for future use via the “–– save_trimmed” parameter and stored in the *trimgalore* directory.

Each fastq sequencing file is separately aligned to both the chosen experimental (“–– experimental”) and spike-in (“––spikein”) genomes in a parallel fashion, with these intermediate alignments available in the *aligned* directory. Misaligned and duplicated reads are removed using the Samtools^26^ and Picard^27^ software suites, with post-filtering alignments and metrics data stored in the *deduplicated* directory. Summaries of these filtering metrics can be found in the *picard_summaries* directory.

Next, the percentages of overlapping reads (independently aligning to both genomes) and spike-in reads (as a fraction of all aligned reads) are calculated for each sample, with the results collected in a single csv file (*overlap_report.csv*). Since the accuracy of normalization can be difficult when the percentage of spike-in reads is too low, the pipeline is designed to automatically exclude any samples with a calculated percentage of spike-in reads below 0.5%. This functionality can be overridden, forcing the pipeline to attempt normalization for all samples, by setting the “––override_spikeinfail” parameter to true. Conversely, PerCell normalization can be disabled (e.g. for analysis of ChIP-seq data lacking spike-in) by setting the parameter “––skip_downsample” to true.

If PerCell normalization is enabled, scaling factors are calculated for each sample (*scaling_factors.csv*) based on the relative number of properly aligned spike-in reads. These factors are used to randomly (via the “––seed” parameter) downsample reads, with the now normalized alignment files stored in the *downsampled* directory. Each immunoprecipitated sample is matched with its corresponding control, and peaks are called using a customized script invoking subcommands from MACS2^28^. This was necessary to avoid MACS2’s default callpeak command and its built-in normalization (based on library size) from obscuring our own (based on spike-in). Each sample’s immunoprecipitated/control pairing is used to score potential peaks using the selected method (“macs2_peak_method”) and stored in the *macs2_subcommand* directory along with narrowPeak files containing peaks passing the cutoff given in the “––macs2_cutoff” parameter. Bedgraphs for each pair of immunoprecipitated and control samples are next compared using MACS2’s bdgcmp command based on the “–– macs2_bigwig_method” parameter, before conversion into the visualizable bigWig format using tools from the UCSC software suite^29^.

Optionally, setting “––skip_idr” to false will use the IDR framework^30^ to identify statistically consistent peaks across two replicates (output in the *idr* directory) based on the chosen “–– idr_cutoff” value. As an alternative, e.g. when a more stringent MACS2 peak calling cutoff is used, consensus peaks can instead be identified using BEDTools^31^ by setting “–– skip_consensus” to false. Finally, the HOMER software suite^32^ can be used to annotate peaksets via the “––skip_annotation” and identify enriched motifs via “––skip_motif” parameters, with the output found within their respective subdirectories inside the *homer* directory.

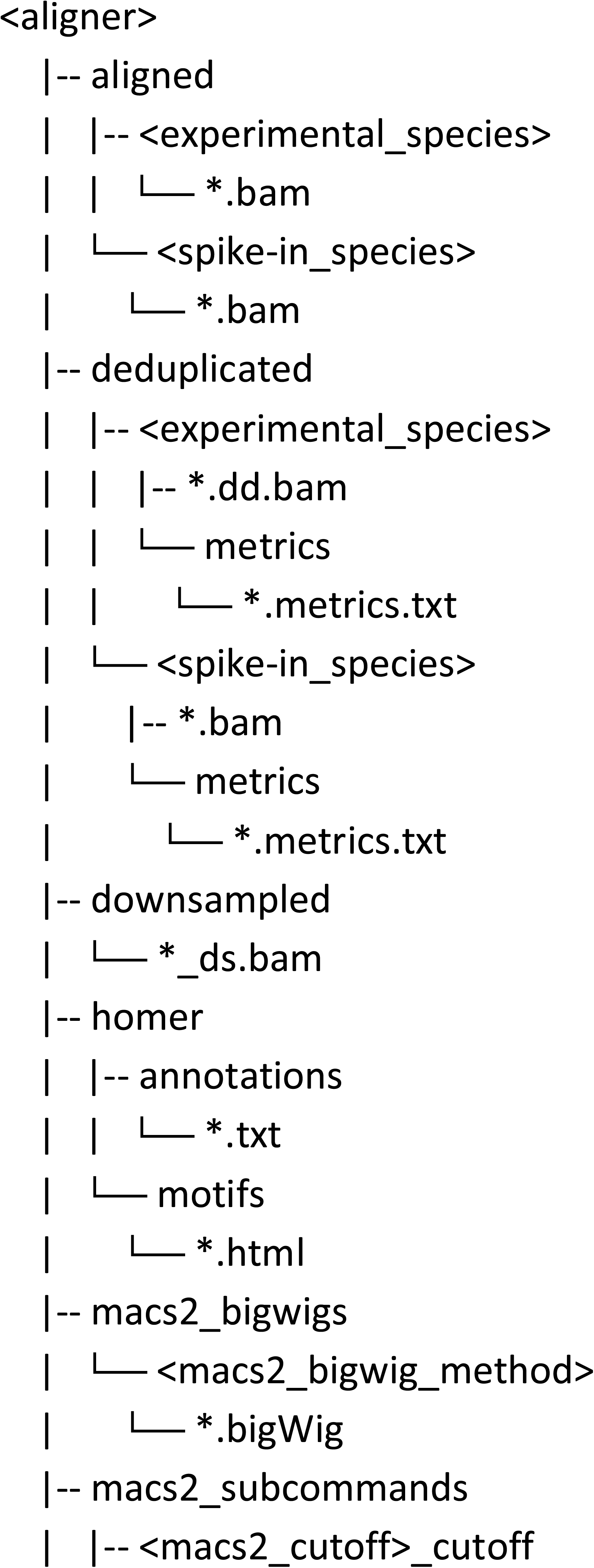
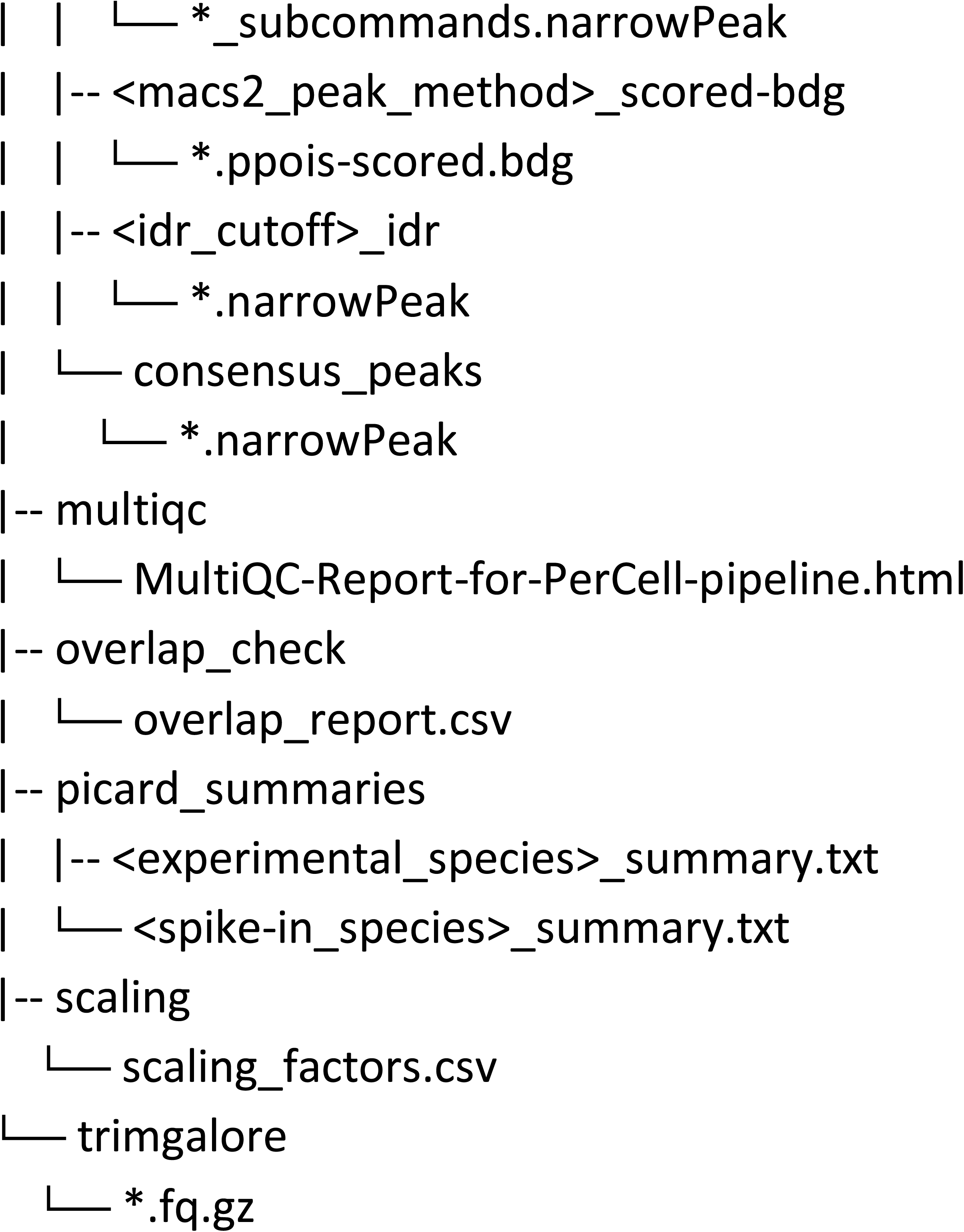

## Troubleshooting

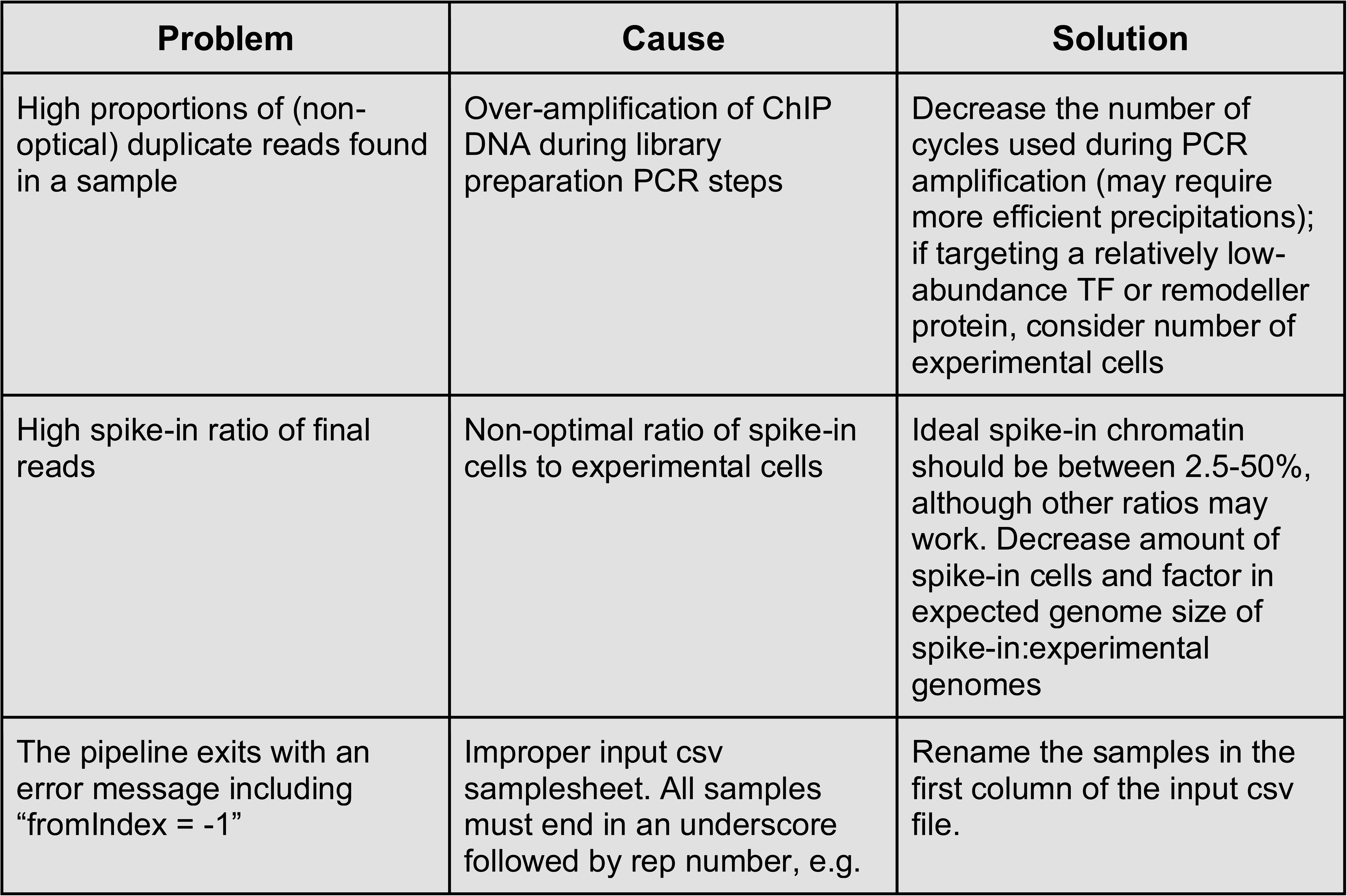
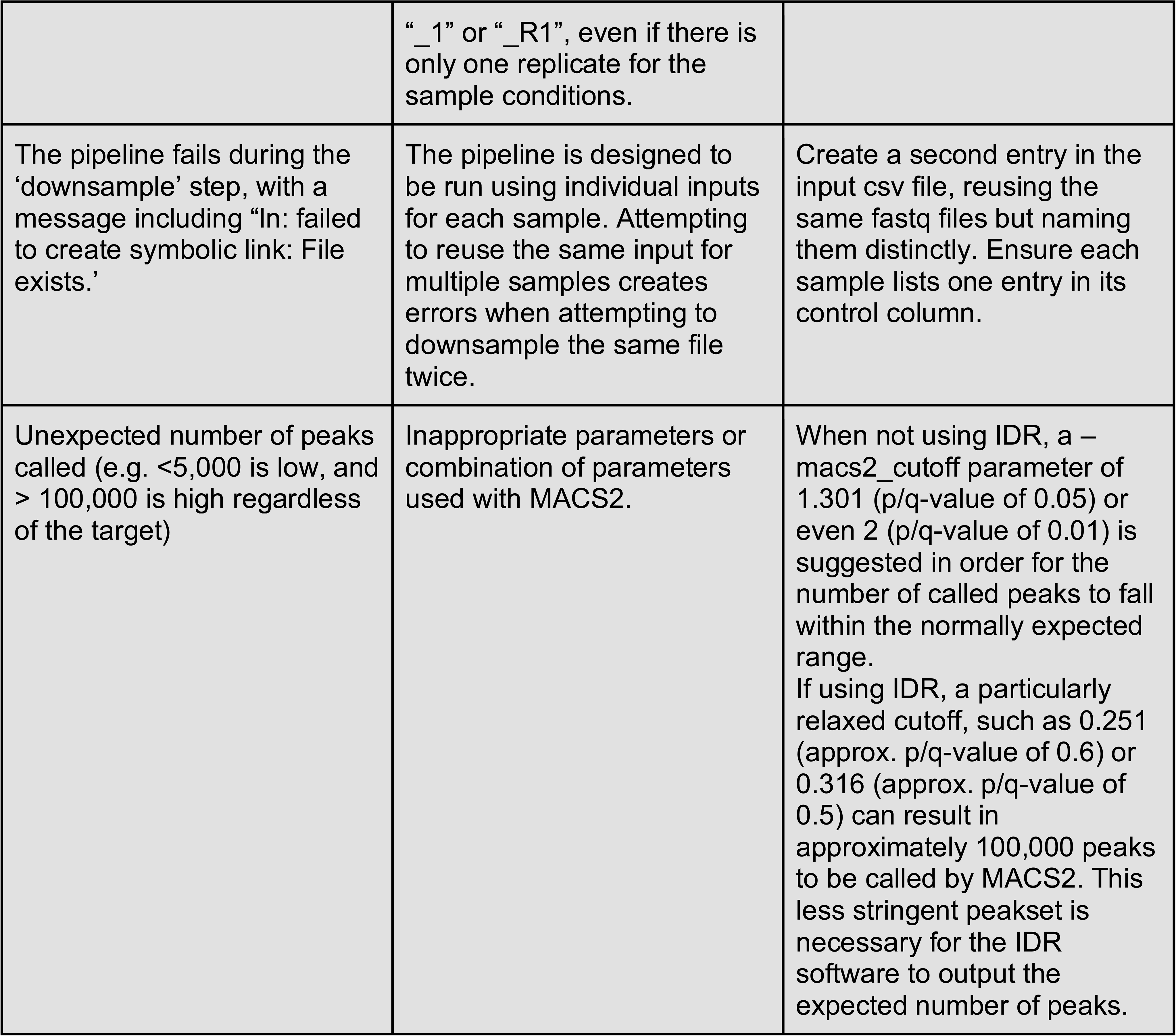

| Problem | Cause | Solution |
| --- | --- | --- |
| High proportions of (non-optical) duplicate reads found in a sample | Over-amplification of ChIP DNA during library preparation PCR steps | Decrease the number of cycles used during PCR amplification (may require more efficient precipitations); if targeting a relatively low-abundance TF or remodeller protein, consider number of experimental cells |
| High spike-in ratio of final reads | Non-optimal ratio of spike-in cells to experimental cells | Ideal spike-in chromatin should be between 2.5-50%, although other ratios may work. Decrease amount of spike-in cells and factor in expected genome size of spike-in:experimental genomes |
| The pipeline exits with an error message including "fromIndex = -1" | Improper input csv samplesheet. All samples must end in an underscore followed by rep number, e.g. | Rename the samples in the first column of the input csv file. |
|  | “_1” or “_R1”, even if there is only one replicate for the sample conditions. |  |
| The pipeline fails during the ‘downsample’ step, with a message including “ln: failed to create symbolic link: File exists.” | The pipeline is designed to be run using individual inputs for each sample. Attempting to reuse the same input for multiple samples creates errors when attempting to downsample the same file twice. | Create a second entry in the input csv file, reusing the same fastq files but naming them distinctly. Ensure each sample lists one entry in its control column. |
| Unexpected number of peaks called (e.g. <5,000 is low, and > 100,000 is high regardless of the target) | Inappropriate parameters or combination of parameters used with MACS2. | When not using IDR, a –macs2_cutoff parameter of 1.301 (p/q-value of 0.05) or even 2 (p/q-value of 0.01) is suggested in order for the number of called peaks to fall within the normally expected range.<br>If using IDR, a particularly relaxed cutoff, such as 0.251 (approx. p/q-value of 0.6) or 0.316 (approx. p/q-value of 0.5) can result in approximately 100,000 peaks to be called by MACS2. This less stringent peakset is necessary for the IDR software to output the expected number of peaks. |

**Supplementary Table S1.** Overlap results verifying pipeline distinguishes between species. : **human (hg38) and mouse (mm39)**. Data are quantified for treatments of DMSO (control), A485 (histone acetyltransferase inhibitor), or Entinostat (histone deacetylase inhibitor), Treated for 8 hours RH4 (Human) with internal cellular Spike-in with C2C12 (Mouse) cells.

| Sample | Experimental Reads | Spike-in Reads | Common Reads | Percent in Common |
| --- | --- | --- | --- | --- |
| DMSO Input Rep1 | 521039 | 148043 | 2306 | 0.345% |
| DMSO Input Rep2 | 1957974 | 471960 | 4231 | 0.174% |
| A485 0.5μM Input Rep1 | 1297108 | 355727 | 2282 | 0.138% |
| A485 0.5μM Input Rep2 | 2888534 | 781066 | 2899 | 0.079% |
| Entinostat 0.5μM Input Rep1 | 969931 | 204427 | 1358 | 0.116% |
| Entinostat 0.5μM Input Rep2 | 2652490 | 898055 | 2953 | 0.083% |
| DMSO H3K27ac Rep1 | 1041982 | 231242 | 2336 | 0.183% |
| DMSO H3K27ac Rep2 | 3410819 | 558869 | 6798 | 0.171% |
| A485 0.5μM H3K27ac Rep1 | 962376 | 242491 | 2194 | 0.182% |
| A485 0.5μM H3K27ac Rep2 | 3180494 | 710326 | 6654 | 0.171% |
| Entinostat 0.5μM H3K27ac Rep1 | 1041186 | 168581 | 1861 | 0.154% |
| Entinostat 0.5μM H3K27ac Rep2 | 2244751 | 409592 | 3438 | 0.130% |
| **Average** | **1847390** | **431698** | **3276** | **0.160%** |

**Supplementary Table S2.** Quantified species read overlap results verifying that the analysis pipeline distinguishes between zebrafish (danRer11) and human (hg38). Data are quantified for control (GFP) injection or PAX3::FOXO1 injected Embryos (Zebrafish) with human cellular spike-in RH30 cells.

| **Sample** | **Experimental Reads** | **Spike-in Reads** | **Common Reads** | **Percent in Common** |
| --- | --- | --- | --- | --- |
| Ctrl-injected H3K27ac Rep1 | 9401727 | 37094793 | 656 | 0.0014% |
| Ctrl-injected H3K27ac Rep2 | 7471932 | 36862309 | 651 | 0.0015% |
| P3F-injected Input Rep1 | 5831485 | 26352450 | 308 | 0.0010% |
| Ctrl-injected Input Rep1 | 6927524 | 26754697 | 349 | 0.0010% |
| P3F-injected Input Rep2 | 5938201 | 25962405 | 397 | 0.0012% |
| Ctrl-injected Input Rep2 | 4985020 | 28366055 | 226 | 0.0007% |
| P3F-injected H3K27ac Rep1 | 7386605 | 32177107 | 594 | 0.0015% |
| P3F-injected H3K27ac Rep2 | 6988345 | 29343096 | 389 | 0.0011% |
| **Average** | **6866355** | **30364114** | **446** | **0.0012%** |

**Supplementary Figure S1.**
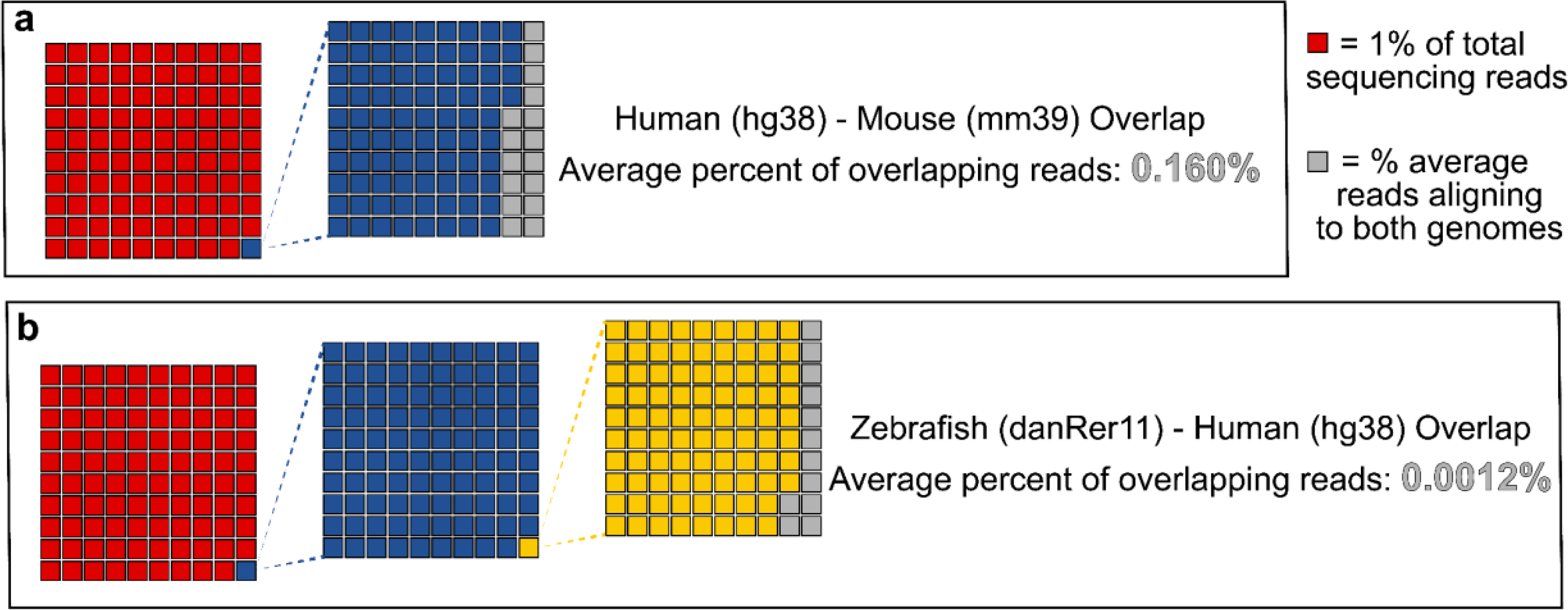
**Graphical representation of minimal fraction of reads aligning to both experimental and spike-in genomes**. **(a)** Human and mouse overlap corresponding to data in Table S1. **(b)** Zebrafish and human species read overlap corresponding to data in Table S2.

**Supplementary Table S3.** Anticipated memory/resource usage for each process in Nextflow-based PerCell analysis pipeline. Note: Not all processes will be required for every run of the pipeline.

| Process | Execution time | Memory usage (RAM) |
| --- | --- | --- |
| INPUT_CHECK:SAMPLESHEET_CHECK | <5s | 5-12 MB |
| CUSTOM_GETCHROMSIZES | 5-20s | 6 MB |
| whitelist_experimental | 5-20s | 6-14 MB |
| whitelist_spikein | 5-20s | 6-14 MB |
| FASTQC | ~5m | ~1 GB |
| TRIMGALORE | 20-30m | ~3 GB |
| BOWTIE2_SPIKEIN | 1m - 1h | 1-4 GB |
| BOWTIE2_EXPERIMENTAL | 1m - 1h | 1-4 GB |
| samtools_spikein | 10s - 8m | 150 MB - 6 GB |
| samtools_experimental | 10s - 8m | 150 MB - 6 GB |
| PICARD_MERGE_SPIKEIN | 5s - 4m | 5 MB - 1.5 GB |
| PICARD_DEDUP_SPIKEIN | 20s - 5m | 3-7 GB |
| PICARD_MERGE_EXPERIMENTAL | 5s - 4m | 5 MB - 1.5 GB |
| PICARD_DEDUP_EXPERIMENTAL | 20s - 5m | 3-7 GB |
| overlap_check | 10s - 8m | 100 MB - 5 GB |
| flagstat | 5s - 1m | 6 MB |
| MULTIQC | 1-30s | 80-100 MB |
| picard_stats | 5-30s | 3 MB |
| calculate | 15-30s | 10-70 MB |
| downsample | 10s - 2m | 10-12 MB |
| macs2_peakcalling | 1m30s - 20m | 150 MB - 3 GB |
| MACS2_CALLPEAK | 15s - 3m | 10 MB - 1 GB |
| homer_findMotifsGenome | 5-45m | 1-2 GB |
| macs2_bdgcmp | 3-25m | 400 MB - 2.5 GB |
| FRIP_SCORE | 30s - 3m | 20 - 300 MB |
| idr | 5s - 2m | 100 MB - 1 GB |

## Data Availability

All ChIP-seq datasets generated with this protocol and presented herein will be deposited in the NCBI Gene Expression Omnibus (GEO, http://www.ncbi.nlm.nih.gov/geo/) following peer-review.

## Code Availability

All code for the PerCell analysis pipeline is publicly available on GitHub (https://github.com/lextallan/PerCell). Downloading the pipeline is not necessary before running it, as Nextflow will automatically download it before the first execution (https://www.nextflow.io/docs/latest/sharing.html#how-it-works).

## Acknowledgements

We thank the Nationwide Children’s Hospital (NCH) Animal Resources Core for their exceptional zebrafish husbandry, especially the Zebrafish Facility team members Dr. Laurie Goodchild, Dr. Carmen Arsuaga, Dr. Lindsey Ferguson, Logan Fehrenbach, Alex Kramer, and Logan Bern. Additional support was provided by the NCH Institute for Genomic Medicine, the NCH Genomics Services Laboratory, and the NCH High-Performance Computing group for assistance in maintaining and using the NCH cluster. We also thank Dr. Matthew Kent for assistance with cloning.

B.Z.S. is grateful to American Cancer Society (RSG-23-1021178-01-DMC), St. Baldrick’s Foundation (Career Development Award), National Institutes of Health (R01GM144601, 1R01HL166520 – 01A1), and intramural funding from Nationwide Children’s Hospital for supporting this work. We are grateful to all members of Stanton and Kendall groups for helpful discussions. G.C.K is grateful for support from an NIH/NCI R01 grant R01CA272872, an Alex’s Lemonade Stand Foundation “A” Award, a V Foundation for Cancer Research V Scholar Award, a CancerFree Kids New Idea Award, and Startup Funds from The Abigail Wexner Research Institute at Nationwide Children’s Hospital. B.Z.S and G.C.K were supported by a Nationwide Children’s Hospital Seed Fund from the Center for Childhood Cancer Research. J.K. is supported by a T32 CA269052 Training Program in Basic and Translational Pediatric Oncology Research predoctoral fellowship. The Institute for Genomic Medicine is funded by the Nationwide Foundation Pediatric Innovation Fund and the Ohio State University Comprehensive Cancer Center grant P30 CA016058. The funders had no role in study design, data collection and analysis, decision to publish, or preparation of the manuscript. Further, the content is solely the responsibility of the authors and does not necessarily represent the official views of the National Institutes of Health.

