## Supplementary figures and images for "Highly quantitative measurement of differential protein-genome binding with PerCell chromatin sequencing"

### Supplementary Figure S1

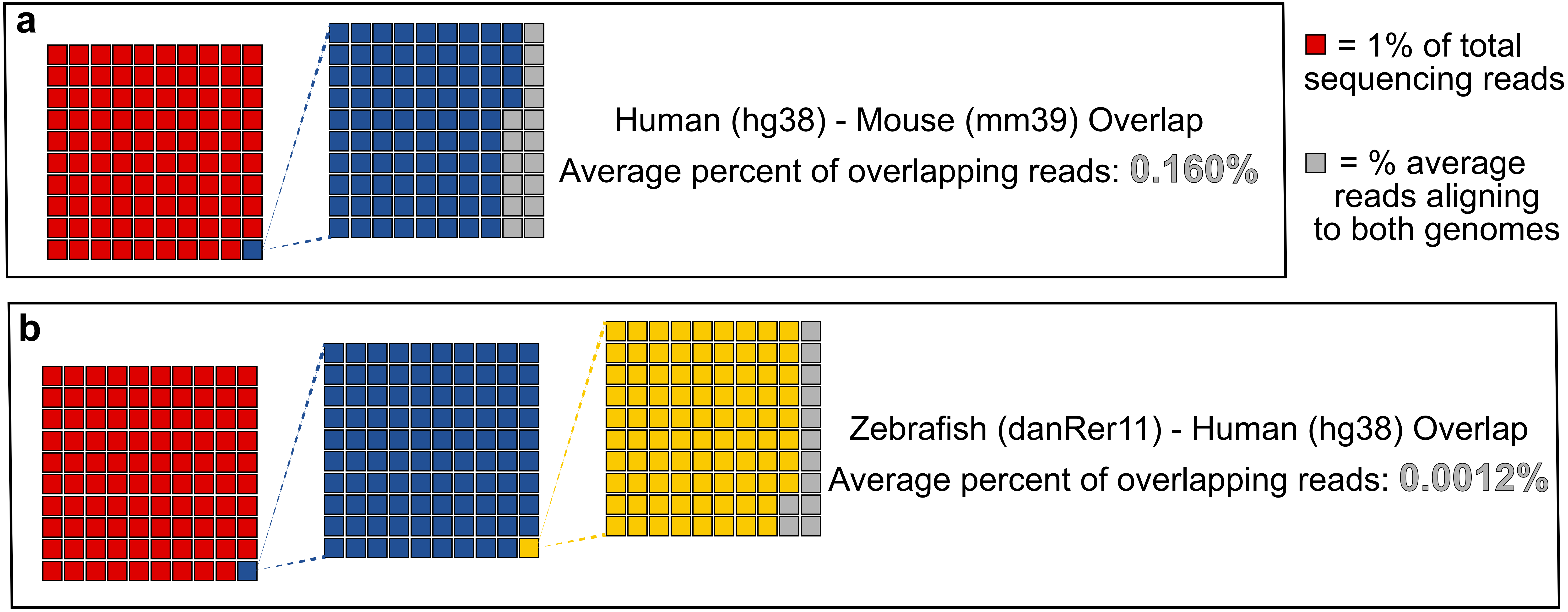
